# Parental Influence on Children’s Educational Achievements: Analysing Direct and Indirect Genetic Effects through Trio-GCTA

**DOI:** 10.64898/2025.12.03.691991

**Authors:** Qiyuan Peng, Espen M. Eilertsen, Rosa Cheesman, Cornelius A. Rietveld, Eivind Ystrom, Alexandra Havdahl

## Abstract

Educational achievement is a key predictor of later-life outcomes, including financial security, social mobility, health, and mortality. Knowing its familial determinants, such as genetic predispositions, is crucial for understanding intergenerational educational mobility and addressing educational inequality. Here, we disentangle direct and indirect genetic effects on educational achievement using trio genome-wide complex trait analysis (Trio-GCTA). Leveraging up to 23,200 genotyped parent–offspring trios from the Norwegian Mother, Father, and Child Cohort Study (MoBa), we find that direct genetic effects explain 15-21% of the variance in educational achievements, including 8th-grade national assessments in Math, Reading, and English, and 10th-grade grade point average (GPA). Notably, indirect genetic effects additionally explain 7-15% of the variance. Both mothers and fathers exert notable influences, exceeding those suggested in previous studies. Moreover, positive gene–environment correlations between parents and offspring show that the intrafamilial environment and children’s genetic predisposition are dependent and mutually reinforcing.

## Introduction

Educational achievement refers to an individual’s performance in school and is closely linked to key psychological, social, economic, and health outcomes across the life course^1^. While children’s learning primarily occurs in educational settings, parents keep playing a crucial role in shaping academic outcomes. Studies have put forth a strong and positive relationship between parental involvement and children’s academic achievement^2^. Additionally, across various societies where data are available, children from less-educated families or those with lower socioeconomic status face greater challenges in achieving comparable levels of academic and social success to their peers from higher-educated or more advantaged backgrounds^3,4^. Understanding the mechanisms behind the intergenerational transmission of educational achievement is crucial for designing interventions to reduce inequalities and support the long-term achievement, health, and well-being of individual children, their families, and society.

The *First Law* of behavioural genetics states that virtually all human traits are heritable to some extent^5,6^, and educational achievement is no exception. Twin studies have consistently supported a prominent role of genetic influences in individual differences in children’s educational achievements, with heritability, which is defined as the proportion of phenotypic variance among individuals that can be attributed to genetic differences within a certain population, estimated at around 60%^7–11^. Heritability estimates based on single-nucleotide polymorphisms (SNPs) are roughly half that level, typically around 30% for educational traits^8,10^. A difference is expected, since SNP-based heritability captures only additive effects tagged by measured or imputed common genetic variants identified through genotyping arrays, excluding contributions from rare variants^12^. By contrast, twin heritability estimates capture the aggregate effects of all inherited DNA differences across the genome within the sample.

Individual differences in educational achievement arise not only from *direct genetic effects*, which reflect the influence of an individual’s own alleles on their own outcomes, but also from *indirect genetic effects*, which occur when alleles in one individual, such as a parent, influence the outcomes of another individual, such as their offspring. Parental genes exert such indirect, environmentally mediated effects on their offspring’s educational outcomes through genetically influenced parental traits, a phenomenon referred to as *genetic nurture*. Using relatedness disequilibrium regression (RDR), Young et al.^13^ estimated that parental genetic nurture effects accounted for 6.6% of the variance in adult educational attainment. However, the total variance explained by parental genetic nurture has not yet been quantified for children’s educational achievement. Evidence from studies using polygenic indices (PGIs) suggests that such parental genetic nurture effects are also present for educational outcomes in childhood, although PGIs capture only a fraction of the underlying genetic signal tagged by measured SNPs. For example, Wang et al.^14^ reported robust evidence for a small but consistent genetic nurture effect on children’s educational outcomes across studies using PGIs (standardised regression coefficient *β* = 0.08, 95% CI [0.07, 0.09]). Importantly, they found that maternal and paternal indirect genetic effects were of comparable magnitude, underscoring the importance of including fathers in educational research. Using Dutch data, Trindade Pons et al.^15^ also identified genetic nurture effects on education, with tentative evidence for more substantial maternal effects. Some studies have found that these effects are partly explained by family socioeconomic status^16–18^, consistent with adoption study evidence showing that adoptive parents with higher income tend to have children with higher educational attainment^19^. Other studies have reported additional mediating effects of parental cognitive performance^20^, maternal health during pregnancy^18^, and parenting behaviours^21^. Although genetic nurture effects exist and various aspects of the family environment have been identified as potential mediators of educational outcomes, such effects are typically modest in magnitude and cannot account for the majority of observed variation in educational achievement. Moreover, current evidence remains inconclusive regarding systematic differences between maternal and paternal genetic nurture effects.

Indirect genetic effects necessarily operate through environmental pathways. Because parents share half of their (autosomal) genetic variants with their children, the genomes of parents and offspring are correlated, giving rise to *gene–environment correlation*. Thus, observed parent–offspring associations may be wholly or partly driven by shared genetic factors^22,23^. For example, parents who transmit to their children a genetic predisposition for higher educational achievement may also create home environments that promote such outcomes, that is, characterised by greater cognitive stimulation, more responsive parenting, and safer, more orderly surroundings. This would produce a positive association between children’s genetic propensity for educational achievement and the quality of their family environment. Additionally, children inheriting genetic tendencies linked to high academic performance may elicit reduced parental investment, such as a “hands-off” approach, leading to a negative gene–environment correlation, which could result in a lower estimate of heritability if this gene-environment correlation is not accounted for. Such gene–environment correlations vitiate any attempt to quantify the importance of family environments without a joint consideration of genetic effects. A review by Jami et al.^24^ reported that gene–environment correlations are pervasive across multiple domains, including depression, criminal behaviour, educational attainment, and substance use in children.

Direct and indirect genetic effects and gene-environment correlations are complex and intertwined. Plomin and Daniels^25^ argued that the myriad idiosyncratic events constituting the ‘environment’ in twin and family studies, while collectively of great importance, are so indefinite and elusive that attempting to study them is a Sisyphean task and thus ‘a gloomy prospect’. These could include parental heritable traits, nurturing offspring, and academic performance, but these are unrelated to parental academic attainment, and hence would be beyond the scope of any design utilising genetic or phenotypic parental educational measures. However, in combination with comprehensive parent-offspring data, the recent development of trio genome-wide complex trait analysis (Trio-GCTA)^26^ allows for disentangling the intertwined genetic and environmental contributions to children’s educational achievement. Trio-GCTA is an extension of genome-wide complex trait analysis (GCTA)^27,28^, a statistical method that estimates heritability based on SNPs across the genome. Using genotyped data from mothers, fathers, and offspring, Trio-GCTA can partition the variance in educational achievement into offspring direct genetic effects, maternal and paternal indirect genetic effects, and gene-environment correlation.

Trio-GCTA offers several methodological advantages. First, it eliminates the risk of reverse confounding, in which an observed association between an exposure and an outcome partly reflects the influence of the outcome on the exposure, a limitation that is common in observational studies^26^. In the Trio-GCTA framework, maternal-, paternal-, and offspring-driven effects are derived from genomic data and cannot be explained by reverse causation, as a child’s educational achievement cannot alter the DNA sequence of their parents. Second, unlike trait-based approaches that rely on specific polygenic indices of measured parental or offspring phenotypes, variance-component methods such as Trio-GCTA can estimate the total contribution of indirect genetic effects without requiring direct measurement of all relevant traits.

In this study, we apply Trio-GCTA to a large sample from the Norwegian Mother, Father and Child Cohort Study (MoBa) to estimate the direct genetic effects of children’s own genotypes and the indirect genetic effects of both mothers and fathers on children’s educational achievement, across multiple educational stages and domains.

## Results

We evaluated the intrafamilial influence on offspring’s educational achievement in 8^th^ grade, as measured by national tests in Math, Reading, and English, as well as grade point average (GPA) at the end of 10^th^ grade, using SNP data from mother-father-offspring trios. Table 1 summarises the results from the Trio-GCTA approach for each measure. Parameter estimates are standardised so that the total variance after conditioning on covariates equals one. Figure 1 shows the variance decomposition for different educational outcomes with parameter estimates from the models with the lowest Akaike’s Information Criteria (AIC) values.

**Figure 1.**
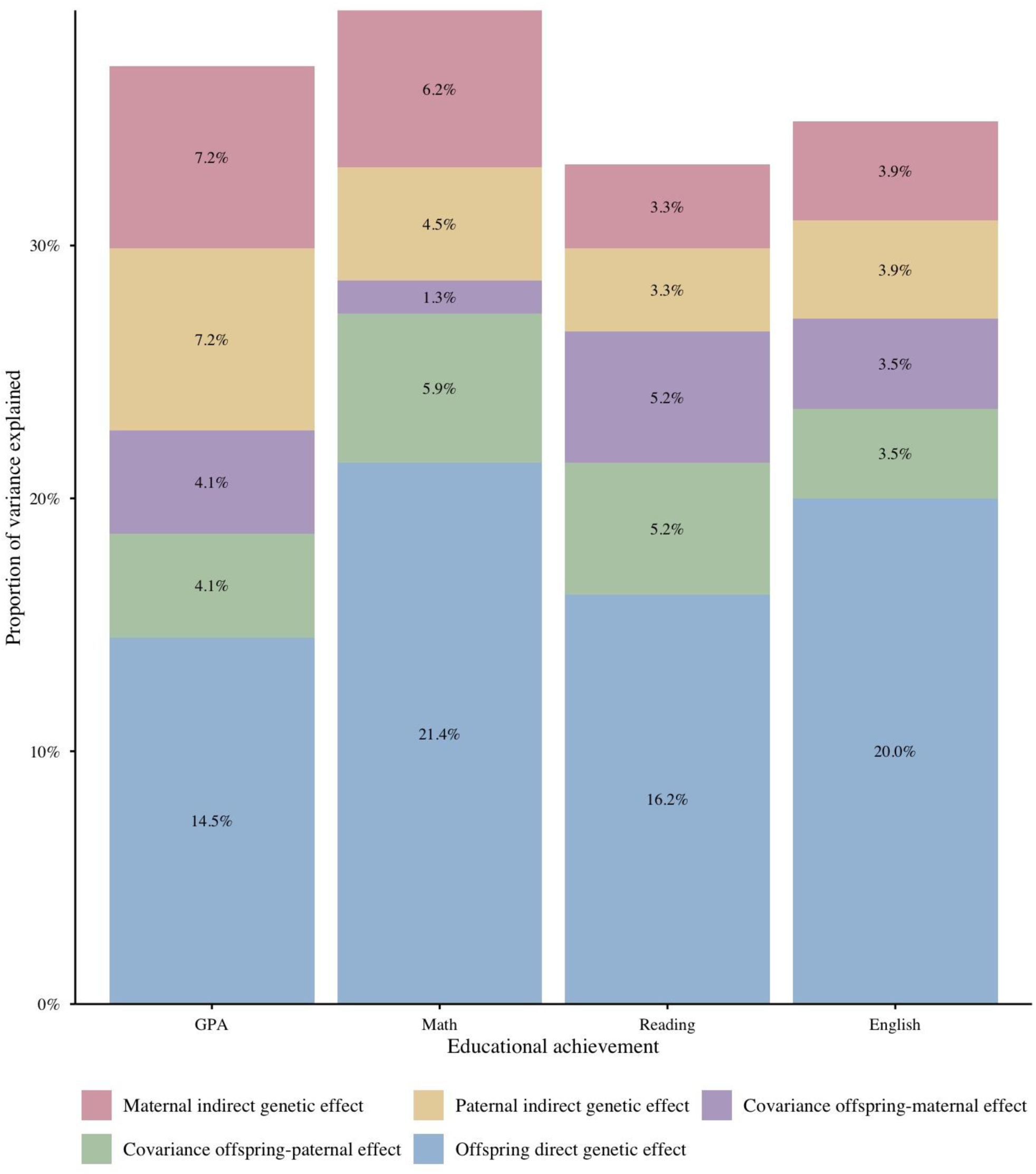
Estimates of direct and indirect genetic components across educational outcomes

**Table 1.**
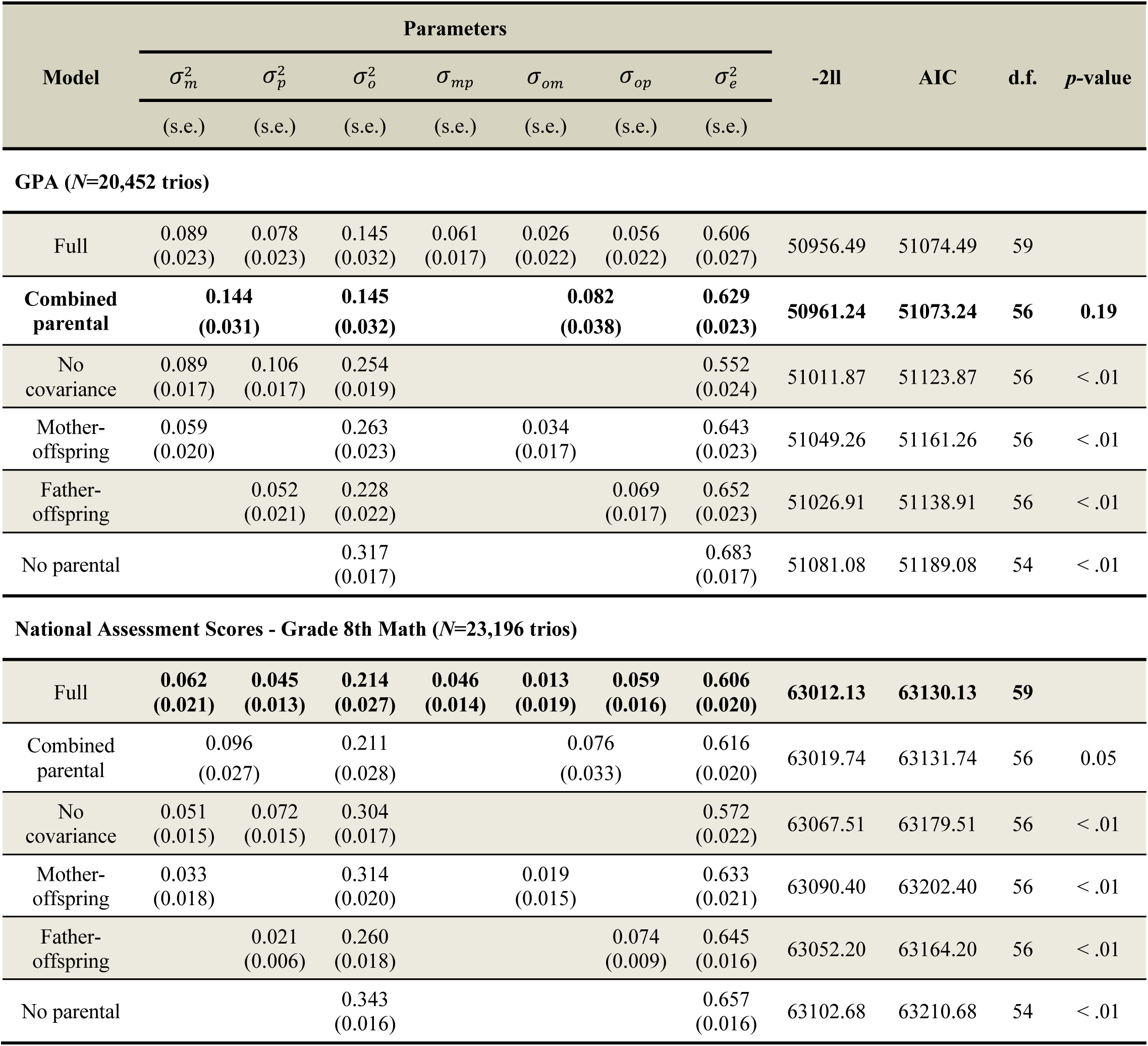
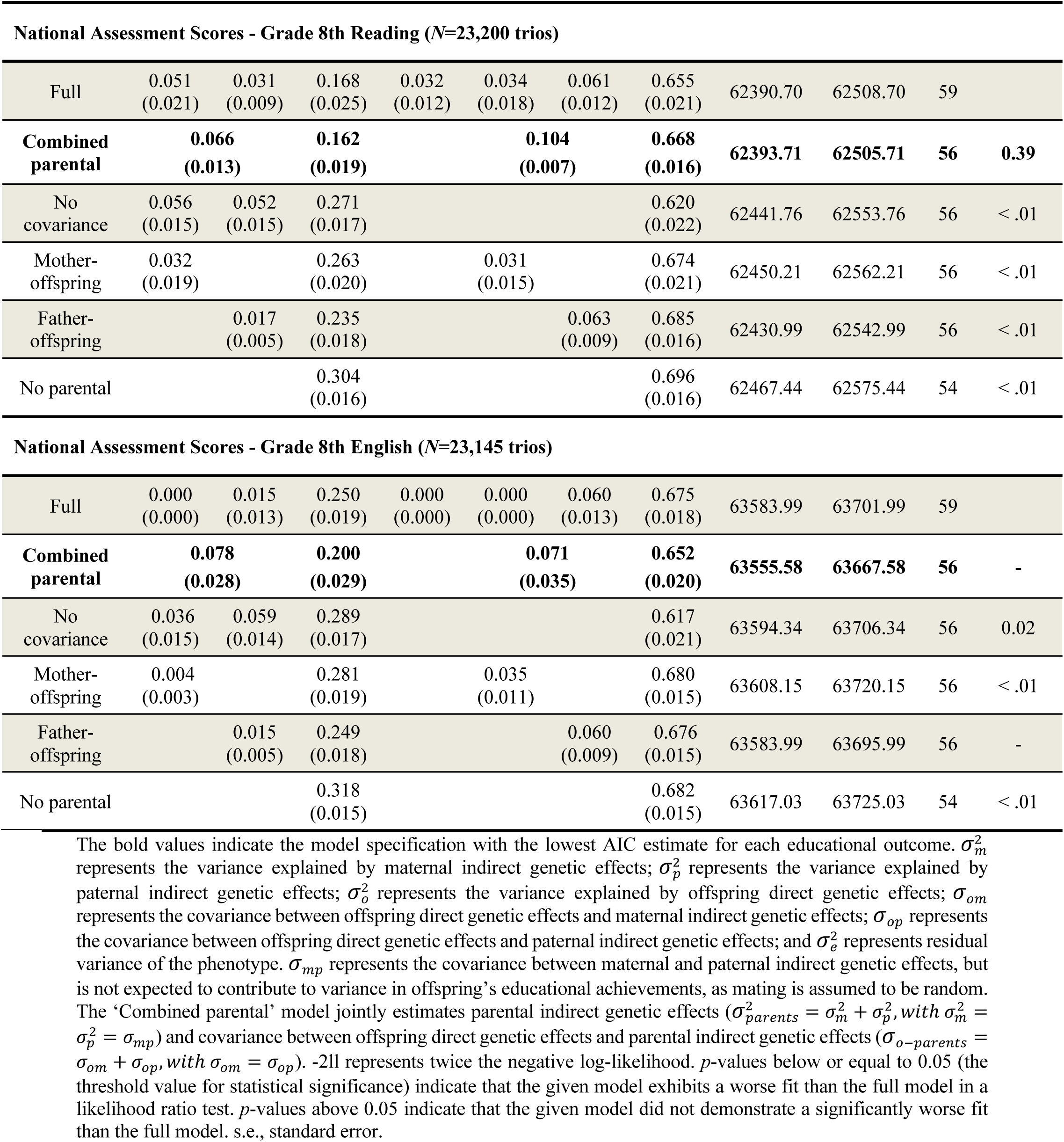
Parameter estimates and fit statistics for each model specification.

The AIC statistics, an index of parsimony, indicate that all the models involving indirect genetic effects provide a (substantial) improvement over the ‘No parental’ model, which only contains the direct genetic effect from offspring. The ‘No covariance’ model, ‘Mother-offspring’ model, and ‘Father-offspring’ model are all nested in the ‘Full’ model, and their fit is much worse than the ‘Full’ model. The ‘Combined parental’ model is also nested in the ‘Full’ model, but is always favoured over the more elaborate ‘Full’ model, except for Math at 8^th^ grade, where the ‘Full’ model provided the best fit.

However, differences in AIC values between the ‘Combined parental’ model and ‘Full’ model were quite small. As such, there is no strong evidence favouring either of these genetic models over the other. Therefore, we discuss the results from both models, characterise the total contribution of indirect genetic effects, and compare the absolute contributions of maternal and paternal indirect genetic effects when relevant. The likelihood ratio tests generally suggested a similar pattern of model fit as the AIC values. This is presented in the *p*-value column of Table 1. At the 5% level, there is no significant loss of fit when considering the ‘Combined parental’ models for GPA and Reading. At the 10% level, there is a significant loss of fit when considering the ‘Combined parental’ models for Math. However, across all GPAs, Math, and Reading outcomes, there is a significant loss of fit compared to the ‘Full’ model by considering the ‘No covariance’ model, ‘Mother-offspring’ model, ‘Father-offspring’ model, and ‘No parental’ model. For English outcomes in 8th grade, we focus on results from the ‘Combined parental’ model, which showed genuine convergence and provided a more stable fit for our dataset. Further details and discussions of the model fitting for English outcomes are presented in the *Supplementary Information*.

Considering the models with the lowest AIC values for GPA, Math, and Reading, and the valid ‘Combined parental’ model for English, the variance explained was 37.1%, 39.4%, 33.2%, and 34.8%, respectively. The variance explained includes components attributable to direct and indirect genetic effects as well as covariances, and it is of interest to inspect the individual components of variance.

Direct genetic effects from offspring accounted for 14.5% (*SE*=3.2%) of the variance in GPA, as well as 21.4% (*SE*=2.7%) of the variance in Math scores, 16.2% (*SE*=1.9%) in Reading scores, and 20.0% (*SE*=2.9%) in English scores. These figures are within-family heritability estimates rather than conventional heritability estimates. In comparison, the combined indirect genetic effects from both parents accounted for 14.4% (*SE*=3.1%) of the variance in GPA, 6.6% (*SE*=1.3%) in Reading, and 7.8% (*SE*=2.8%) in English. For Math, the maternal indirect genetic effect accounted for 6.2% (*SE*=2.1%) of the variance, and the paternal indirect genetic effect accounted for 4.5% (*SE*=1.3%), for a total of 10.7%.

The covariance between direct and indirect genetic effects was always positive for all educational achievements, explaining 8.2% (*SE*=3.8%) of the variance for GPA, 10.4% (*SE*=0.7%) for Reading, and 7.1% (*SE*=3.5%) for English. For Math, the covariance between direct and maternal indirect genetic effects explained 1.3% of the variance (*SE*=1.9%), and the covariance between direct and paternal indirect genetic effects explained 5.9% of the variance (*SE*=1.6%). All of these findings correspond to a positive gene–environment correlation, suggesting that partially overlapping sets of genes may be involved in both direct and indirect genetic effects. To some extent, these effects accompany each other and jointly contribute to increased phenotypic variance. Table 2 presents the gene–environment correlations among genetic effects across various educational outcomes.

**Table 2.**
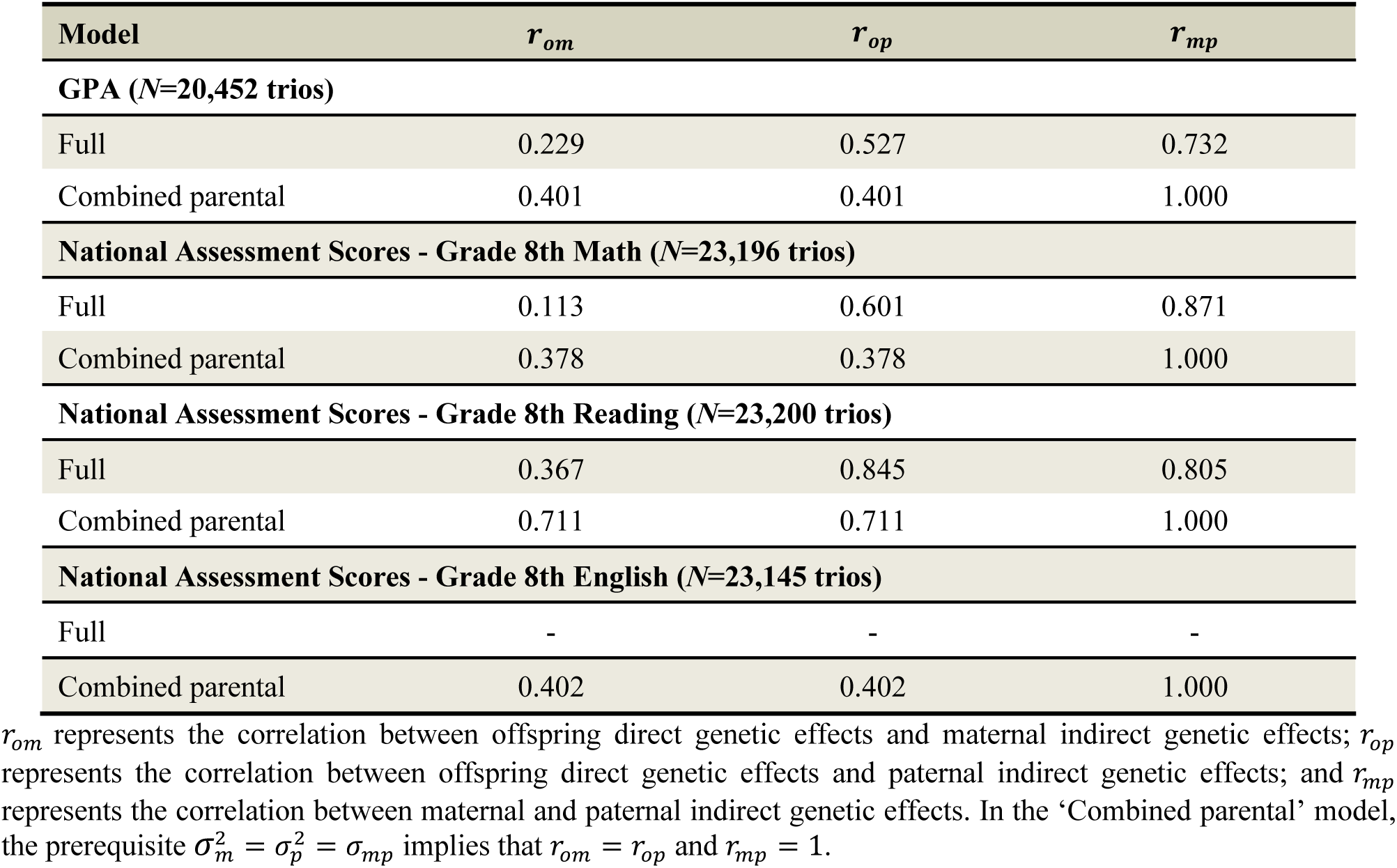
Gene–environment correlations between genetic effects across educational outcomes.

| Model | $r_{om}$ | $r_{op}$ | $r_{mp}$ |
| --- | --- | --- | --- |
| **GPA (N=20,452 trios)** |  |  |  |
| Full | 0.229 | 0.527 | 0.732 |
| Combined parental | 0.401 | 0.401 | 1.000 |
| **National Assessment Scores - Grade 8th Math (N=23,196 trios)** |  |  |  |
| Full | 0.113 | 0.601 | 0.871 |
| Combined parental | 0.378 | 0.378 | 1.000 |
| **National Assessment Scores - Grade 8th Reading (N=23,200 trios)** |  |  |  |
| Full | 0.367 | 0.845 | 0.805 |
| Combined parental | 0.711 | 0.711 | 1.000 |
| **National Assessment Scores - Grade 8th English (N=23,145 trios)** |  |  |  |
| Full | - | - | - |
| Combined parental | 0.402 | 0.402 | 1.000 |

$r_{om}$ represents the correlation between offspring direct genetic effects and maternal indirect genetic effects; $r_{op}$ represents the correlation between offspring direct genetic effects and paternal indirect genetic effects; and $r_{mp}$ represents the correlation between maternal and paternal indirect genetic effects. In the ‘Combined parental’ model, the prerequisite $\sigma_m^2 = \sigma_p^2 = \sigma_{mp}$ implies that $r_{om} = r_{op}$ and $r_{mp} = 1$ .

## Discussion

Using the Trio-GCTA approach in a large-scale sample of up to 23,200 Norwegian parent–offspring trios, we find consistent evidence that both offspring direct genetic effects and parental indirect genetic effects contribute to offspring educational achievement across multiple stages of schooling. Given the small model-fitting statistics, we find no strong evidence favouring either the ‘Full’ or ‘Combined parental’ model over the other in the current dataset. Nevertheless, it remains essential to account for both maternal and paternal indirect genetic effects, regardless of whether they are assumed to differ or be equivalent, when examining the variance in offspring educational achievement. Notably, we observed positive covariance between direct and indirect genetic effects across all outcomes and model specifications. Collectively, these findings underscore the importance of accounting for intra-familial genetic influences when studying the developmental origins of educational achievement.

Direct genetic effects, typically assumed to be the predominant component of heritability estimates, were robustly evident across all educational outcomes in our Trio-GCTA analyses. In both the ’Full’ and ’Combined parental’ models, direct genetic effects accounted for 15% to 21% of the total variance for standardised test scores in Math, Reading, and English at 8^th^ grade, and for GPA at the end of 10^th^ grade. These estimates are consistent with previous findings based on family-based molecular genomic designs, such as Relatedness Disequilibrium Regression (RDR) estimates^13^, which estimated the heritability of educational attainment to be 17% and can be viewed as a simplified version of the Trio-GCTA approach using the midparent genotype rather than separating maternal and paternal effects. Both Trio-GCTA and RDR approaches can provide precise estimates of heritability with negligible bias due to the environment. Our estimates of direct genetic effects are lower than SNP-based heritability estimates derived from single-generation unrelated individuals (e.g., GREML), which typically report ∼30% heritability for educational traits^8,10^. Notably, this ∼30% value is very close to the total variance explained by the combined direct and indirect effects in our analysis. This is expected, as GREML estimates cannot fully remove the upward bias due to indirect genetic effects from relatives even after excluding close relatives^13,29^. Young et al.^13^ suggested that GREML-SNP estimates can be interpreted as estimates of the variance explained by the combined direct and indirect effects of transmitted alleles, rather than heritability. In contrast, our study allows for the disentanglement of direct genetic effects on offspring educational achievements from parental indirect genetic effects, thereby providing a more accurate estimate of direct genetic influence on offspring outcomes.

Unlike trait-based approaches, our model does not specify the parental phenotypes underlying the indirect genetic effects and thus remains agnostic to the interactive processes between parents and offspring. Instead, it partitions the aggregated genetic variance into components attributable to maternal, paternal, and offspring genetic variation, along with their covariances. The direct genetic effect captures the combined influence of all genotyped SNPs in the offspring, regardless of whether these effects are mediated through multiple interactions with parental behaviours. Likewise, the indirect genetic effects capture the combined influence of all genotyped SNPs in mothers and fathers, which may ultimately represent a cascade of interactions with offspring behaviours. Although the specific environmental pathways or mechanisms underlying these indirect genetic effects cannot be identified, we can nonetheless draw broader inferences from the observed variance components.

Indirect maternal genetic effects accounted for 3% to 9% of the variance across different educational achievements in both the ‘Full’ and ‘Combined parental’ models, while indirect paternal genetic effects explained between 3% and 8%. These estimates indicate that parental genetic variants exert measurable influences on offspring’s educational achievement through environmentally mediated pathways rather than through genetic transmission alone. In other words, maternal and paternal genotypes appear to contribute to variability in offspring outcomes via the rearing environment shaped by parents’ genetically influenced traits and behaviours. Thus, our results provide empirical support to theoretical models emphasising the importance of parental characteristics and family environment in shaping children’s educational achievement across developmental stages.

In the ‘Combined parental’ models, parental indirect genetic effects explained 14.4% of the variance in GPA, 9.6% of the variance in Math, 6.6% of the variance in Reading, and 7.8% of the variance in English. These results may partly reflect that GPA carries greater weight for future academic progression and opportunities for both children and their parents, motivating stronger parental engagement and environmental support.

Furthermore, our results suggest that both mothers and fathers play a more important role in shaping offspring’s educational achievement than has been indicated in previous empirical studies. Our results also highlight the need for greater caution when assuming parental symmetry in genetic studies of the intergenerational transmission of education. Previous behavioural research has shown that the association between parental involvement and children’s educational outcomes is equally strong for fathers and mothers^14,30,31^. However, our results from the ‘Full’ model indicate that maternal and paternal genetic nurture effects are not equivalent, particularly for mathematics achievement. In conjunction with their differing covariances with direct genetic effects, this suggests that the differences may reflect not only variation in magnitude but also in underlying mechanisms. Moreover, both the ‘Full’ and ‘Combined parental’ models consistently demonstrate that genetic differences among parents are associated with phenotypic differences in their offspring, irrespective of the underlying mechanism. Taken together, renewed emphasis on the roles of both parents is warranted. Wherever possible, both mothers and fathers should be included in research and intervention efforts.

Our estimates of indirect parental genetic effects are derived without assuming that all parental influences are shared among siblings. Plomin^32^ argued that family experiences may not be shared equally among siblings but may instead be specific to each individual^25^. Trio-GCTA provides a direct test of this hypothesis, as our estimates rely solely on detecting effects of parental genotypes on offspring traits after accounting for the direct effects of the offspring’s own genotype. However, our estimates are limited to parental effects captured by the additive effects of genotyped SNPs, which likely underestimate the full additive genetic contribution^12^. In addition, the total indirect parental effect may include genetic effects from rare variants, non-additive genetic influences, or de novo mutations, and may also correlate with broader social environmental factors. As such, our Trio-GCTA estimates represent a conservative, lower-bound approximation of the total indirect parental effect.

All covariances involving both direct and indirect genetic effects were positive, indicating that correlations between these effects were consistently positive across all educational achievements. In other words, we observed the positive gene–environment correlations here. This suggests that the intrafamilial environment is not independent of children’s own genetic liability for educational achievement. The implication for family environment research is that genetic effects may confound any naïve association with specific aspects of the family, meaning such associations should not be interpreted as purely causal social effects.

Moreover, because the positive gene–environment correlations between offspring and either mothers or fathers were well below 1, only part of the same genetic variation is shared between direct and indirect genetic effects. This indicates that parents influence children’s education primarily through environmental pathways other than those directly related to traits associated with their own educational achievement. Notably, in the ‘Full’ model across different educational outcomes, the gene–environment correlations between mothers and offspring were much smaller than those between fathers and offspring, yet the variances explained by maternal and paternal indirect genetic effects were similar. This pattern suggests that mothers and fathers may affect children’s educational outcomes through partly different environmental pathways, which is also reflected in the correlations between maternal and paternal indirect genetic effects (<1).

In ecology and animal studies^33–39^, there has been extensive discussion of the correlation between direct and indirect parental genetic effects. The evolutionary dynamics of a trait can be strongly shaped by the correlation—or genetic covariance—between parental indirect genetic effects and offspring direct genetic effects. Empirical evidence indicates that this association may be either positive or negative. A negative correlation may help maintain genetic variation across generations but will slow or even reverse the rate at which selection improves a trait^35^. In contrast, a positive correlation implies that direct and indirect parental genetic effects act in the same direction, which can rapidly accelerate microevolutionary change^35^.

These insights can be applied to our context. Normally, we do not think of the ‘environment’ as something that can evolve, unless it is determined, at least in part, by maternally or paternally expressed genes^33,36^. For a heritable trait, if parental indirect genetic effects exist, then the family environment shaped by such genes can evolve alongside offspring direct genetic effects. This co-evolution will influence how traits under selection change over time^36,39^. In the presence of a positive correlation, parental indirect and offspring direct genetic effects act together, increasing the observed variance in offspring educational achievement and potentially accelerating the evolution of education-related behaviours. However, to date, parameter estimates for such correlations remain scarce in both human behavioural research and studies of wild populations, making it difficult to determine the conditions that produce correlations of different directions or magnitudes^36^.

There are several limitations that should be kept in mind when interpreting the results of our study. First, the differences between the competing models concerning model fit statistics (AIC and likelihood values) were generally small. Therefore, the statistical support for choosing any specific model is not strong. We mainly demonstrate the presence of offspring direct genetic effects and parental indirect genetic effects, showing that these sources of variability are interconnected. Larger sample sizes are needed to quantify these estimates more accurately. Second, our analyses were restricted to children with both parents and a Norwegian context, and the study was based on genotype data of European ancestry. These sample restrictions may limit the generalizability of our findings beyond this group. Moreover, the family structure in Trio-GCTA includes only one child per family (mother-father-child), whereas most Norwegian families have more than one child. Indirect genetic effects from siblings could also contribute to individual differences and may correlate with parental effects. Third, estimates of indirect genetic effects may be affected by assortative mating and population stratification, as shown in previous polygenic indices studies of educational outcomes^40,41^. When assortative mating occurs for a heritable trait, this results in increased trait-specific genetic and phenotypic variances in the child generation, and it would induce covariance between the non-transmitted alleles of the mother and the transmitted alleles of the father^42–44^. Such processes could, in principle, inflate estimates of indirect genetic effects. Population stratification may also contribute to bias if allele frequencies correlate with social or regional environmental factors that also influence educational outcomes. Nevertheless, robustness checks (see *Supplementary Information*) indicated that our results were rather insensitive to excluding genetically more related individuals. Future work should aim to estimate parental effects while accounting for assortative mating and population. Finally, although the possible influence of selection bias in MoBa cannot be ruled out, attrition in our target outcomes is likely minimal because educational data were obtained from compulsory, standardised national tests and administrative registries^45,46^. Acknowledging these limitations, our findings remain consistent with the view that parents play a substantial role, both genetically and environmentally, in shaping their children’s educational achievements. Continued advances in family-based genomic designs and larger, more diverse samples will enable even stronger inferences about the mechanisms underlying intergenerational educational inequality.

## Conclusion

Investigating genetic contributions to educational achievement from a within-family framework offers valuable insights into the intertwined genetic and environmental pathways that shape children’s educational achievement. Using the MoBa cohort, we identified and quantified both direct and indirect genetic effects on educational achievement for Math, Reading, and English at 8th grade, and for GPA at the end of 10th grade. These findings highlight the importance of considering the roles of both mothers and fathers in shaping children’s educational outcomes. Our results underscore the importance of accounting for intrafamilial influences, particularly parental indirect genetic effects and gene–environment correlations, when studying the development of educational outcomes. The observed positive gene–environment correlations imply that direct and indirect genetic effects to some extent accompany each other, increasing the variability in children’s educational outcomes. Crucially, failing to account for indirect parental genetic effects risks biasing the identification of genetic variants associated with children’s educational outcomes. Overall, this study demonstrates the utility of genomic family-based designs and the Trio-GCTA method for disentangling the direct and indirect genetic effects on children’s educational achievements.

## Methods

### The Norwegian Education System

Norway is a large country with relatively few inhabitants. The Norwegian welfare system is well-developed and based on universal entitlements^47^. Compulsory education in Norway begins at age six and lasts for ten years, covering primary education (grades 1–7) and lower secondary education (grades 8–10)^48,49^. School intake is generally determined by residential location, and only a small percentage of students attend private schools. Primary and lower secondary education follows the principle of a unified school system that aims to provide equal and individually adapted education to all students. There is a common national curriculum for primary and secondary education; however, within this framework, municipal and county authorities, schools, and teachers can influence the practical implementation of teaching and training.

Students who complete primary and lower secondary education are entitled to up to four years of upper secondary education and training. Counties are responsible for upper secondary education and training, as well as post-secondary vocational education. In higher education, the degree structure aligns with the Bologna Process, comprising primarily a three-year bachelor’s degree, a two-year master’s degree, and a three-year PhD program. With the exception of a few private university colleges, all higher education institutions are publicly owned, and the national government is responsible for overseeing higher education.

### Data, Measures and Sample

#### Data Sources

The data used was via the project SUBPU, which is approved by the Regional Committees for Medical and Health Research Ethics (ref. 2017/2205). The University of Oslo is responsible for the data handling in SUBPU and has conducted a Data Protection Impact Assessment (DPIA) in collaboration with the Norwegian Agency for Shared Services in Education and Research (Sikt; ref. 962088).

##### Statistics Norway (SSB)

Statistics Norway is the national statistical institute of Norway and the main producer of official statistics. They are responsible for collecting, producing, and communicating statistics related to the economy, population, and society at national, regional, and local levels. Statistics Norway also conducts extensive research and analysis activities. This study used the microdata database of population and education from Statistics Norway.

##### Norwegian Mother, Father, and Child Cohort Study (MoBa)

The Norwegian Mother, Father, and Child Cohort Study (MoBa) is a prospective population-based pregnancy cohort study conducted by the Norwegian Institute of Public Health. Pregnant women were recruited from across Norway from 1999 to 2008 (*N* = 112,908 recruited pregnancies)^50,51^. The women consented to initial participation in 41% of the pregnancies. The total cohort includes approximately 114,500 children, 95,200 mothers and 75,200 fathers^50^. Blood samples were obtained from both parents during pregnancy and from mothers and children (umbilical cord) at birth^52^. The current study is based on version 12 of the quality-assured data files released for research in January 2019. The establishment of MoBa and initial data collection was based on a license from the Norwegian Data Protection Agency and approval from the Regional Committees for Medical and Health Research Ethics (REK). The MoBa cohort is currently regulated by the Norwegian Health Registry Act.

Genotyping of MoBa has been conducted through multiple research projects (HARVEST, SELECTIONpreDISPOSED, and NORMENT), spanning several years; using varying selection criteria, genotyping arrays, and genotyping centres^51^. A standardised pipeline was used for pre-imputation quality control (QC), phasing, imputation, and post-imputation QC of the genotyping data. Details of the MoBa genotyping efforts and the MoBaPsychGen QC pipeline are described elsewhere^51^.

#### Measures

##### Grade Point Average (GPA)

Data on grade point average (GPA) were obtained from Norway’s National Education Database (NUDB), managed by Statistics Norway (SSB).

GPA is the final assessment score reflecting students’ academic performance upon completion of compulsory education at the end of lower secondary school (grade 10, approximately age 16). At the end of grade 10, students are required to take a centrally administered written examination in one of three subjects: Norwegian or Sami, Mathematics, or English. Additionally, most students are required to take one locally administered oral examination. The oral examination may cover any school subject. For subjects without a written or oral examination, final grades are based on teachers’ overall assessment of students’ achievement throughout the academic year. All grades are given numerically on a scale from 1 (lowest) to 6 (highest). GPA is calculated by averaging these overall assessment and examination grades, rounding the result to two decimal places, and then multiplying by 10. For analytical purposes, we standardised GPA within each academic year to have a mean of zero and a standard deviation of one.

Historical GPA records in the database date back to 2001. Some students may lack GPA records due to various exemptions, including recent immigration status, enrollment in individualised education programs, recent transfers between schools, or exemptions requested by parents. Additionally, students exhibiting high absenteeism, academic misconduct (e.g., cheating), expulsion due to disorderly conduct, or failure to participate in exams also do not receive GPA scores. Moreover, students with valid grades in fewer than half of their subjects are not assigned an official GPA and are excluded from national statistics. Consequently, approximately 90% of Norwegian students have valid GPA data recorded.

Because GPA from lower secondary school is used as a critical criterion for selection into upper secondary education, students likely perceive the GPA received in grade 10 as significantly higher stakes compared to grades from previous years. Consequently, GPA serves as an important predictor of a student’s future educational trajectory and employment opportunities, making it a particularly relevant outcome to examine.

##### National Assessment Scores

National assessment scores for Mathematics, Reading, and English in grade 8 were obtained through linkage with Norway’s National Education Database (NUDB), maintained by Statistics Norway (SSB).

The national assessment in grade 8, administered digitally every autumn since 2007, is the first nationwide test encountered by students (approximately age 13) after entering lower secondary school. According to regulations, all students are required to participate in these national tests, with exemptions granted to students with documented special education needs and those following introductory language courses. Consequently, approximately 96% of all Norwegian students have valid national assessment data available.

Assessment results are reported to students on a five-level scale, with cut-offs at the 10th, 30th, 70th, and 90th percentiles. For analytical purposes, we used the raw scores and standardised each test score within subject and year to have a mean of zero and a standard deviation of one.

The primary purpose of the national assessment is to monitor educational outcomes at the school, municipal, and national levels, facilitating the development of improvement strategies both locally and centrally, rather than evaluating individual students. Therefore, these assessments are generally considered low stakes for students.

##### Selection of Parent-Offspring Trios

After post-imputation quality control of genotype data, 44,017 full mother-father-offspring trios (with one complete trio in the parent generation) were retained^51^. We excluded the latest version of withdrawal participants. We further selected parent-offspring trios with children’s educational achievement data collected in grade 8 and grade 10. In families with multiple genotyped children having educational data, we selected the oldest child.

Our analysis relies on identifying different types of genetic effects based on genetic relatedness within and between parent-offspring trios. To do so, we computed a genomic relatedness matrix (GRM)^28^, which empirically estimates the genetic similarity among all individuals in the sample. Because closely related individuals can disproportionately influence genetic variance estimates and introduce confounding due to shared environmental factors not specified in the model^12^, we applied a threshold of 0.10 for the maximum allowed genetic relatedness between any two individuals (ignoring parent-offspring pairs) in the main analysis. This threshold aligns with previous Trio-GCTA studies^26,53–55^, effectively balancing the exclusion of closely related individuals and maintaining a sufficiently large number of parent-offspring trios. We also applied alternative thresholds of 0.075, 0.050, and 0.025 to select parent-offspring trios for robustness checks reported in the *Supplementary Information*. The GRM and the selection of individuals were computed using the “bottom-up” algorithm implemented through functions from the OpenMendel project^56^.

Consequently, the final analytic sample included 20,542 trios with GPA data, 23,196 trios with mathematics data in grade 8, 23,200 trios with Norwegian reading data in grade 8, and 23,145 trios with English data in grade 8.

### Statistical Analysis

Trio-GCTA^26^ is applied in the present study to estimate the direct and indirect genetic effects underlying children’s educational achievement. Trio-GCTA is extended from GCTA^28^ and M-GCTA^57^ to quantify the importance of direct and indirect genetic effects within the nuclear family by using parent-offspring trios. The direct and indirect genetic effects here represent the combined influence of all genetic variants tagged by genotyped and imputed SNPs. The model can be formulated as a multiple linear regression model for the offspring’s phenotype

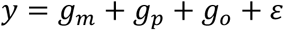

𝑔*_o_* represents the direct genetic effects from offspring, 𝑔*_m_* and 𝑔*_p_* represent the indirect genetic effects from mother and father, and 𝜀 represents the residual effects from all other sources of variability. Both genetic and residual effects are assumed to follow a multivariate normal distribution. The different genetic effects are assumed to be correlated across individuals according to their genomic relatedness matrix values. Furthermore, it is assumed that random mating occurs in the population. Therefore, the total variance decomposition of the offspring’s phenotype is

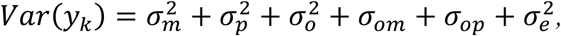

where 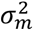 and 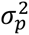 corresponds to variance attributable to indirect maternal and paternal genetic effects, respectively, whereas 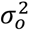 is the variance due to direct genetic effects. The components 𝜎*_om_* and 𝜎*_op_* are the covariances between the direct offspring genetic effect and the indirect maternal and paternal genetic effects, respectively. The covariance terms are necessary if partially the same genes contribute directly and indirectly. These parameters quantify the extent to which the same variants contribute to direct and indirect genetic effects. Because parents and offspring are related, a positive covariance between direct and indirect genetic effects will increase variability in the phenotype, whereas a negative covariance will decrease variability. With respect to the offspring, the maternal and paternal genetic effects form part of the environment, so these covariance terms may therefore also be interpreted as measuring variability due to gene-environment correlations. The component 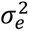 is the residual variance of the phenotype. The residual variance estimate may include genetic effects not captured by SNPs included in the analysis, unique environmental effects, and shared environmental effects not captured by SNPs. The covariance between indirect maternal and paternal genetic effects 𝜎*_mp_* is estimated, but is not expected to contribute to the variance in offspring’s educational achievement, as mating is assumed to be random.

We fitted five more alternative, simpler models of how children’s phenotype may depend on genetic effects, nested within the general model described above (**‘Full’ model**). Firstly, it could be sufficient to consider that mother and father contribute the same to the offspring’s phenotype. This is obtained by substituting the indirect maternal and paternal genetic effects with a combined indirect parental genetic effect (𝑔*_mp_* = 𝑔*_m_* + 𝑔*_p_*). Young et al.^13^ used to model that maternal and paternal genetic effects are equally important and perfectly correlated (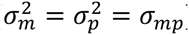), and equally correlated with the direct genetic effect (𝜎*_om_* = 𝜎*_op_*) when they introduced the relatedness disequilibrium regression (RDR) method. Then the variance accounted for by the combined parental indirect genetic effects 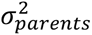 is equal to 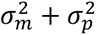, and the covariance with direct genetic effect 𝜎*_o_*_-*parents*,_ is equal to 𝜎*_om_* + 𝜎*_op_*. We refer to this as the ‘**Combined parental**’ model. Then, the next model, which is named as the ‘**No covariance**’ model, estimated fewer parameters, dropping either covariance parameters for the direct and indirect genetic effects. Eaves et al.^57^ proposed M-GCTA for jointly estimating the variance explained by direct genetic effects, indirect maternal genetic effects and their covariance concerning an offspring phenotype. This ‘**Mother-offspring**’ model can be obtained with the constraints 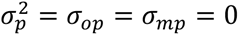. Similarly, for only considering direct genetic effects, indirect paternal genetic effect and their covariance on children’s educational achievement, we set the constraints 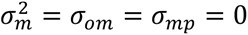 for the ‘**Father-offspring**’ model. We also considered a model without any indirect genetic effects of parents (‘**No parental**’ model). This is equivalent to the genomic-relatedness-matrix restricted maximum likelihood method, implemented in GCTA software^28^. Specific variance components estimated in each model tested in the Trio-GCTA are shown in Table 3.

**Table 3.** Models and variance components estimated in each model.

| Model | Variance decomposition |
| --- | --- |
| Full | $\sigma_m^2 + \sigma_p^2 + \sigma_o^2 + \sigma_{om} + \sigma_{op} + \sigma_e^2$ |
| Combined parental | $\sigma_{parents}^2 + \sigma_o^2 + \sigma_{o-parents} + \sigma_e^2$ |
| No covariance | $\sigma_m^2 + \sigma_p^2 + \sigma_o^2 + \sigma_e^2$ |
| Mother-offspring | $\sigma_m^2 + \sigma_o^2 + \sigma_{om} + \sigma_e^2$ |
| Father-offspring | $\sigma_p^2 + \sigma_o^2 + \sigma_{op} + \sigma_e^2$ |
| No parental | $\sigma_o^2 + \sigma_e^2$ |

$\sigma_m^2$ represents the variance explained by maternal indirect genetic effects; $\sigma_p^2$ represents the variance explained by paternal indirect genetic effects; $\sigma_o^2$ represents the variance explained by offspring direct genetic effects; $\sigma_{om}$ represents the covariance between offspring direct genetic effects and maternal indirect genetic effects; $\sigma_{op}$ represents the covariance between offspring direct genetic effects and paternal indirect genetic effects; and $\sigma_e^2$ represents residual variance of the phenotype.

We fitted all these six models to GPA, 8^th^ grade National Assessment scores for Math, Reading and English separately. In all analyses, we additionally included an intercept term and the fixed effects of child sex, genotype batches, imputation batches and principal components of mothers and fathers.

Model fit was assessed using AIC. AIC, an index of parsimony, is efficient at selecting the model closest to the true model when the true model is not present in the candidate models, particularly when the true model is complex and/or contains small effects^58^. We also conducted likelihood ratio tests, comparing the goodness of fit of the full model with that of the nested models for each outcome. However, there are challenges associated with interpreting likelihood ratio tests using family data^59,60^. We are not aware of work examining the interpretation of likelihood ratio tests in the context of GREML methods that involve direct and indirect genetic effects. We therefore relied on AIC for selecting models with the best fit at each outcome. Thus, the model is considered the best-fitting model with the lowest AIC value.

All models were estimated using the Julia programming language^61^, via the package GREMLModels.jl^62^.

## Supporting information

Supplementary Information, and will be used for the link to the file on the preprint site.

## Acknowledgments

The data access and management costs of SUBPU are financed by the Research Council of Norway (RCN), the European Research Council, and the Department of Psychology (UiO). This work was performed on the TSD (Tjenester for Sensitive Data) facilities, owned by the University of Oslo, operated and developed by the TSD service group at the University of Oslo, IT-Department (USIT). The authors would like to acknowledge the work of SUBPU data managers Clara Timpe and Oda van Jole. The computations were performed on resources provided by Sigma2 - the National Infrastructure for High-Performance Computing and Data Storage in Norway (ref. NS9867S). We are grateful to all the participating families in Norway who take part in the MoBa, an ongoing cohort study. We thank the Norwegian Institute of Public Health (NIPH) for generating high-quality genomic data. For generating high-quality genomic data, we thank the Norwegian Institute of Public Health (NIPH), the HARVEST collaboration, the NORMENT Centre at the University of Oslo, the Center for Diabetes Research at the University of Bergen, deCODE Genetics, the Research Council of Norway, the South-Eastern and Western Norway Regional Health Authorities, Stiftelsen KG Jebsen, the Trond Mohn Foundation, and the Novo Nordisk Foundation. This project has received funding from the European Union’s Horizon Europe research and innovation programme under the Marie Skłodowska-Curie grant agreement (ESSGN 101073237). E.M.E., R.C., and E.Y. were supported by the European Research Council (ERC) (GeoGen; Grant agreement No. 101045526). E.Y. was also supported by the Research Council of Norway (288083, 336078, and 331640) and the Swedish Research Council (2024-06499). R.C. was supported by the Jacobs Foundation (2023-1510-00). A.H. was supported by the Research Council of Norway (#336085), the South-Eastern Norway Regional Health Authority (#2020022), and the European Union’s Horizon Europe Research and Innovation programme (FAMILY #101057529). C.A.R. gratefully acknowledge funding from the Dutch Research Council (VI.VIDI.231E.001). The funder had no role in study design, data collection and analysis, preparation of the manuscript, or decision to publish.

## Data availability

Data from the Norwegian Mother, Father and Child Cohort Study is managed by the Norwegian Institute of Public Health and the Educational Registry is managed by Statistics Norway; these data sources can be made equally available to researchers, provided approval from the Regional Committees for Medical and Health Research Ethics, compliance with the EU General Data Protection Regulation (GDPR), and approval from the data owners. The consent given by the participants and the Norwegian Statistics Act does not open for storage of data on an individual level in repositories or journals. Researchers who want access to data sets for replication should apply through helsedata.no. Further details on how to obtain data access can be found at https://www.fhi.no/en/studies/moba/ for MoBa, and at https://www.ssb.no/en/data-til-forskning for Statistics Norway.

## Code availability

The code used in this study is available upon request from the first author. Example code for fitting variance-component models structured according to relationship matrices with GREMLModels.jl is provided at: https://github.com/espenmei/GREMLModels.jl.

## Ethical Considerations

The Norwegian registry and MoBa data used was from the project SUBPU, which is approved by Committees for Medical and Health Research Ethics (#2017/2205). The University of Oslo is responsible for the data handling of SUBPU and has conducted a Data Protection Impact Assessment (DPIA) in collaboration with the Norwegian Agency for Shared Services in Education and Research (Sikt; #962088). SUBPU has agreements with the MoBa and the Statistics Norway for data linkage and usage. MoBa is regulated by the Norwegian Health Registry Act. Informed consent was obtained from all participating mothers and fathers.

## Notes

https://github.com/espenmei/GREMLModels.jl

### Competing Interest Statement

The authors have declared no competing interest.

