## Supplementary Information, and will be used for the link to the file on the preprint site. for "Parental Influence on Children’s Educational Achievements: Analysing Direct and Indirect Genetic Effects through Trio-GCTA"

#### I. Model Convergence and Singular Fit Issues for English Outcomes

For English outcomes in 8th grade, it is difficult to determine the best-fitting model. Although the 'Full' model formally converged, it yielded a boundary solution, with several variance and covariance components estimated as zero (e.g., $\sigma_m^2 = \sigma_{om} = \sigma_{mp} = 0$). This phenomenon is referred to as a singular fit, which typically indicates that the model is overparameterized relative to the available data, resulting in unstable estimates and reduced inferential validity.

While singular models are statistically well defined (i.e., the true maximum likelihood estimate may indeed lie on the boundary of the parameter space), they raise several practical concerns: they are often associated with overfitting and low statistical power, increased risks of numerical instability or mis-convergence, and the potential inappropriateness of standard inferential procedures such as likelihood ratio tests[1]. In such cases, likelihood-based comparisons become unreliable due to boundary estimation or singularities in the covariance matrix.

There is no gold standard for how to deal with singularity and which random-effects specification to choose. Several strategies have been proposed within a frequentist framework. One approach is to avoid fitting overly complex models to ensure that variance–covariance matrices can be estimated with sufficient precision (Matuschek et al., 2017). Another is to apply model selection procedures that balance predictive accuracy against risks of overfitting and inflated type I error[2,3]. Alternatively, the "keep it maximal" principle recommends fitting the most complex model consistent with the experimental design, removing only those random-effects terms necessary to achieve a non-singular fit[4].

Specifically, for the English outcome in our study, the log-likelihood of the 'Full' model was worse than that of the nested 'Combined parental' model, suggesting that the optimiser may have stopped based on numerical convergence criteria without reaching a statistically meaningful optimum. A similar singular fit issue was also observed in the 'Mother-offspring' model for the English outcome. Thus, for English, we focus on the results from the 'Combined parental' model, which is more appropriate for this dataset and exhibits genuine convergence.

**Supplementary Table 1 Parameter estimates and fit statistics for each model specification of English**

| Model | Parameters $\sigma_m^2$ (s.e.) | $\sigma_p^2$ (s.e.) | $\sigma_o^2$ (s.e.) | $\sigma_{mp}$ (s.e.) | $\sigma_{om}$ (s.e.) | $\sigma_{op}$ (s.e.) | $\sigma_e^2$ (s.e.) | -2ll | AIC | d.f. | $p$-value |
|---|---|---|---|---|---|---|---|---|---|---|---|
| **National Assessment Scores - Grade 8th English ($N$=23,145 trios)** | | | | | | | | | | | |
| Full | 0.000 (0.000) | 0.015 (0.013) | 0.250 (0.019) | 0.000 (0.000) | 0.000 (0.000) | 0.060 (0.013) | 0.675 (0.018) | 63583.99 | 63701.99 | 59 | |
| **Combined parental** | **0.078 (0.028)** | | **0.200 (0.029)** | | **0.071 (0.035)** | | **0.652 (0.020)** | **63555.58** | **63667.58** | **56** | **-** |
| No covariance | 0.036 (0.015) | 0.059 (0.014) | 0.289 (0.017) | | | | 0.617 (0.021) | 63594.34 | 63706.34 | 56 | 0.02 |
| Mother-offspring | 0.004 (0.003) | | 0.281 (0.019) | | 0.035 (0.011) | | 0.680 (0.015) | 63608.15 | 63720.15 | 56 | < .01 |
| Father-offspring | | 0.015 (0.005) | 0.249 (0.018) | | | 0.060 (0.009) | 0.676 (0.015) | 63583.99 | 63695.99 | 56 | - |
| No parental | | | 0.318 (0.015) | | | | 0.682 (0.015) | 63617.03 | 63725.03 | 54 | < .01 |

The bold values indicate the model specification with the lowest AIC estimate for English. $\sigma_m^2$ represents the variance explained by maternal indirect genetic effects; $\sigma_p^2$ represents the variance explained by paternal indirect genetic effects; $\sigma_o^2$ represents the variance explained by offspring direct genetic effects; $\sigma_{om}$ represents the covariance between offspring direct genetic effects and maternal indirect genetic effects; $\sigma_{op}$ represents the covariance between offspring direct genetic effects and paternal indirect genetic effects; and $\sigma_e^2$ represents residual variance of the phenotype. $\sigma_{mp}$ represents the covariance between maternal and paternal indirect genetic effects, but is not expected to contribute to variance in offspring's educational achievements, as mating is assumed to be random. The 'Combined parental' model jointly estimates parental indirect genetic effects ($\sigma_{parents}^2 = \sigma_m^2 + \sigma_p^2, with\, \sigma_m^2 = \sigma_p^2 = \sigma_{mp}$) and covariance between offspring direct genetic effects and parental indirect genetic effects ($\sigma_{o-parents} = \sigma_{om} + \sigma_{op}, with\, \sigma_{om} = \sigma_{op}$). -2ll represents twice the negative log-likelihood. $p$-values below or equal to 0.05 (the threshold value for statistical significance) indicate that the given model exhibits a worse fit than the full model in a likelihood ratio test. $p$-values above 0.05 indicate that the given model did not demonstrate a significantly worse fit than the full model. s.e., standard error.

### II. Robustness checks for GPA outcome

**Supplementary Table 1 Parameter estimates and fit statistics for each model specification under different thresholds for the maximum allowed genetic relatedness between any two individuals (GPA)**

| Model for GPA | Parameters | | | | | | | -2ll | AIC | d.f. | *p*-value |
|---|---|---|---|---|---|---|---|---|---|---|---|
| | $\sigma_m^2$ (s.e.) | $\sigma_p^2$ (s.e.) | $\sigma_o^2$ (s.e.) | $\sigma_{mp}$ (s.e.) | $\sigma_{om}$ (s.e.) | $\sigma_{op}$ (s.e.) | $\sigma_e^2$ (s.e.) | | | | |
| **Threshold = 0.1 (*N*=20,452 trios)** | | | | | | | | | | | |
| Full | 0.089 (0.023) | 0.078 (0.023) | 0.145 (0.032) | 0.061 (0.017) | 0.026 (0.022) | 0.056 (0.022) | 0.606 (0.027) | 50956.49 | 51074.49 | 59 | |
| **Combined parental** | **0.144 (0.031)** | | **0.145 (0.032)** | | **0.082 (0.038)** | | **0.629 (0.023)** | **50961.24** | **51073.24** | **56** | **0.19** |
| No covariance | 0.089 (0.017) | 0.106 (0.017) | 0.254 (0.019) | | | | 0.552 (0.024) | 51011.87 | 51123.87 | 56 | < .01 |
| Mother-offspring | 0.059 (0.020) | | 0.263 (0.023) | | 0.034 (0.017) | | 0.643 (0.023) | 51049.26 | 51161.26 | 56 | < .01 |
| Father-offspring | | 0.052 (0.021) | 0.228 (0.022) | | | 0.069 (0.017) | 0.652 (0.023) | 51026.91 | 51138.91 | 56 | < .01 |
| No parental | | | 0.317 (0.017) | | | | 0.683 (0.017) | 51081.08 | 51189.08 | 54 | < .01 |
| **Threshold = 0.075 (*N*=19,074 trios)** | | | | | | | | | | | |
| Full | 0.081 (0.025) | 0.069 (0.025) | 0.166 (0.035) | 0.061 (0.019) | 0.023 (0.024) | 0.050 (0.024) | 0.611 (0.029) | 47310.28 | 47428.28 | 59 | |
| **Combined parental** | **0.136 (0.034)** | | **0.167 (0.034)** | | **0.072 (0.041)** | | **0.625 (0.024)** | **47312.75** | **47424.75** | **56** | **0.48** |
| No covariance | 0.080 (0.018) | 0.094 (0.018) | 0.266 (0.020) | | | | 0.561 (0.025) | 47354.52 | 47466.52 | 56 | < .01 |
| Mother-offspring | 0.050 (0.022) | | 0.270 (0.024) | | 0.035 (0.018) | | 0.645 (0.024) | 47379.56 | 47491.56 | 56 | < .01 |
| Father-offspring | | 0.042 (0.022) | 0.241 (0.024) | | | 0.064 (0.018) | 0.653 (0.024) | 47364.54 | 47476.54 | 56 | < .01 |
| No parental | | | 0.322 (0.018) | | | | 0.678 (0.018) | 47402.59 | 47510.59 | 54 | < .01 |

| | | | | | | | | | | | |
|---|---|---|---|---|---|---|---|---|---|---|---|
| **Threshold = 0.05 (*N*=15,734 trios)** | | | | | | | | | | | |
| Full | 0.070 (0.030) | 0.045 (0.014) | 0.149 (0.037) | 0.049 (0.019) | 0.048 (0.027) | 0.069 (0.018) | 0.619 (0.029) | 38994.35 | 39112.35 | 59 | |
| **Combined parental** | **0.102 (0.021)** | | **0.146 (0.025)** | | **0.122 (0.008)** | | **0.630 (0.022)** | **38995.35** | **39107.35** | **56** | **0.80** |
| No covariance | 0.087 (0.021) | 0.075 (0.022) | 0.281 (0.024) | | | | 0.557 (0.030) | 39033.63 | 39145.63 | 56 | < .01 |
| Mother-offspring | 0.042 (0.027) | | 0.265 (0.029) | | 0.051 (0.022) | | 0.641 (0.030) | 39040.70 | 39152.70 | 56 | < .01 |
| Father-offspring | | 0.020 (0.008) | 0.253 (0.026) | | | 0.072 (0.012) | 0.655 (0.022) | 39037.89 | 39149.89 | 56 | < .01 |
| No parental | | | 0.334 (0.022) | | | | 0.666 (0.022) | 39061.64 | 39169.64 | 54 | < .01 |
| **Threshold = 0.025 (*N*=7,589 trios)** | | | | | | | | | | | |
| Full | 0.033 (0.028) | 0.092 (0.055) | 0.170 (0.058) | 0.038 (0.031) | 0.074 (0.019) | 0.098 (0.041) | 0.533 (0.060) | 19040.99 | 19158.99 | 59 | |
| **Combined parental** | **0.118 (0.041)** | | **0.185 (0.050)** | | **0.148 (0.016)** | | **0.550 (0.046)** | **19043.64** | **19155.64** | **56** | **0.45** |
| No covariance | 0.058 (0.044) | 0.150 (0.044) | 0.336 (0.049) | | | | 0.455 (0.064) | 19052.64 | 19164.64 | 56 | < .01 |
| Mother-offspring | 0.010 (0.011) | | 0.332 (0.056) | | 0.056 (0.029) | | 0.602 (0.045) | 19062.90 | 19174.90 | 56 | < .01 |
| Father-offspring | | 0.079 (0.056) | 0.288 (0.058) | | | 0.088 (0.045) | 0.545 (0.061) | 19050.62 | 19162.62 | 56 | 0.02 |
| No parental | | | 0.395 (0.045) | | | | 0.605 (0.045) | 19066.00 | 19174.00 | 54 | < .01 |

The bold values indicate the model specification with the lowest AIC estimate for GPA. $\sigma_m^2$ represents the variance explained by maternal indirect genetic effects; $\sigma_p^2$ represents the variance explained by paternal indirect genetic effects; $\sigma_o^2$ represents the variance explained by offspring direct genetic effects; $\sigma_{om}$ represents the covariance between offspring direct genetic effects and maternal indirect genetic effects; $\sigma_{op}$ represents the covariance between offspring direct genetic effects and paternal indirect genetic effects; and $\sigma_e^2$ represents residual variance of the phenotype. $\sigma_{mp}$ represents the covariance between maternal and paternal indirect genetic effects, but is not expected to contribute to variance in offspring's educational achievements, as mating is assumed to be random. The 'Combined parental' model jointly estimates parental indirect genetic effects ($\sigma_{parents}^2 = \sigma_m^2 + \sigma_p^2, with\, \sigma_m^2 = \sigma_p^2 = \sigma_{mp}$) and covariance between offspring direct genetic effects and parental indirect genetic effects ($\sigma_{o-parents} = \sigma_{om} + \sigma_{op}, with\, \sigma_{om} = \sigma_{op}$). -2ll represents twice the negative log-likelihood. *p*-values below or equal to 0.05 (the threshold value for statistical significance) indicate that the given model exhibits a worse fit than the full model in a likelihood ratio test. *p*-values above 0.05 indicate that the given model did not demonstrate a significantly worse fit than the full model. s.e., standard error.
